# Olduvai Domains Downregulate Mitochondrial Pathways to Promote Human Brain Evolution and Neoteny

**DOI:** 10.1101/2024.10.21.619278

**Authors:** Jonathon G. Keeney, David Astling, Vanessa Andries, Karl Vandepoele, Nathan Anderson, Jonathan M. Davis, Pamela Lopert, Manisha Patel, Ken Jones, Jonathan Vandenbussche, Kris Gevaert, An Staes, Santos Franco, Natasia Paukovich, Beat Vögeli, Frans van Roy, James M. Sikela

## Abstract

Olduvai (formerly DUF1220) protein domains, encoded by the NBPF gene family, have undergone the greatest human lineage-specific copy-number expansion of any coding sequence in the genome and strongly correlate with brain size and neuron number across primates. Here we show that Olduvai domains act in a dosage-dependent manner to suppress mitochondrial metabolism. Transcriptomic, proteomic, and live-cell imaging analyses of cells overexpressing NBPF1 (which encodes seven Olduvai domains) reveal pronounced downregulation of mitochondrial pathways, including electron transport chain components and NADH dehydrogenase activity, as well as reduced mitochondrial abundance. By limiting energy availability, this suppression delays cellular maturation and developmental timing. We propose that the resulting prolongation of neurogenesis increases neuron production, providing a mechanistic link between Olduvai copy number expansion and the evolutionary enlargement of the human brain. This dosage-sensitive mitochondrial regulation may also contribute to broader neotenic features of human development, offering a unifying molecular mechanism for brain expansion and the neotenic traits that distinguish humans from other primates.

## Introduction

Olduvai protein domains (formerly DUF1220; Sikela and van Roy 2017) have undergone a striking human lineage-specific copy number expansion, representing the largest increase of any coding sequence in the human genome (Fortna et al. 2004; Popesco et al. 2006). Copy numbers rose progressively across Anthropoid primates—first in monkeys, then apes, and most dramatically in humans—adding more than 160 human-specific copies and bringing the total to approximately 300 in modern humans (O’Bleness et al. 2012). These domains are encoded primarily by the *NBPF* gene family (Vandepoele et al. 2005) and fall into six primary clades: three conserved across primates (CON1–3) and three that underwent human-lineage-specific amplification (HLS1–3) (O’Bleness et al. 2012), all of which are intrinsically disordered (Issaian et al. 2019; Paukovich et al. 2022).

Olduvai copy number shows a strong linear correlation with brain size and cortical neuron number across primates, as well as with normal and disease-associated variation in human brain size and cognitive performance (Dumas et al. 2012; Keeney et al. 2015a; Zimmer and Montgomery 2015; Davis et al. 2015a, 2015b, 2019). Higher Olduvai dosage has been shown to increase neural stem cell proliferation (Keeney et al. 2015a), consistent with the observed positive association between Olduvai copy number and cortical neuron number. Previous work raised the possibility that Olduvai contributes to developmental neoteny—a slowing of maturation that prolongs juvenile traits, including extended neurogenesis (Keeney et al. 2014). However, the underlying molecular mechanism remained unclear.

In this study, we investigated Olduvai function by overexpressing *NBPF1* (encoding seven Olduvai domains) in the low endogenous Olduvai human cell line DLD1Tr21. Using integrated transcriptomics, proteomics, and live-cell imaging, we identified consistent downregulation of mitochondrial pathways. This metabolic restraint provides a plausible mechanistic link between Olduvai copy number expansion, the promotion of neoteny, and the evolution of large brains in primates and humans. The finding is particularly noteworthy given recent reports connecting reduced mitochondrial function to developmental slowing and neoteny (Namba et al. 2020; Diaz-Cuadros et al. 2023; Iwata et al. 2023; Liu and Auwerx 2023; Casimir et al. 2024; Lindhout et al. 2024).

## Materials and Methods

### Detailed experimental procedures are provided in the Supplementary Methods

#### Cell culture and live-cell imaging

DLD1Tr21-derived cell lines stably expressing NBPF1-IRES-EGFP or EGFP control (DLD1Tr21/NBPF1 and DLD1Tr21/Mock, respectively) were generated using a Tet-On inducible system as previously described (Vandepoele et al. 2008; Van de Wetering et al. 2002). Expression of *NBPF1* was induced with 2 μg/ml doxycycline (Dox). EGFP was verified by immunofluorescence for transcriptomics and proteomics (Fig S1). For mitochondrial imaging, cells were labeled with 200 nM MitoTracker CMXRos and imaged every 4 h using an IncuCyte live-cell imaging system under standard culture conditions (37°C, 5% CO_2_).

#### RNA sequencing

DLD1Tr21/NBPF1 and DLD1Tr21/Mock cells were cultured with or without Dox for 24 h (four biological replicates per condition). Total RNA was extracted, and poly(A)-selected mRNA was used to prepare cDNA libraries with the Illumina TruSeq RNA Sample Preparation Kit v2. Libraries were sequenced as single-end 100 bp reads on an Illumina HiSeq 2000 platform. On average, 20 million reads were obtained per sample.

#### RNA-seq data analysis

Reads were aligned to the human genome (hg19/GRCh37) using TopHat (v2.0.9) (Kim et al. 2013) and transcripts assembled with Cufflinks (v2.1.1) (Trapnell et al. 2010) guided by RefSeq annotations. Gene-level expression was quantified as FPKM values. Differentially expressed genes were identified using the limma package in R (Smyth 2005) (false discovery rate [FDR] < 0.05, fold-change > 1.5). Gene set enrichment analysis was performed with GSEA (v2.0.13) (Subramanian et al. 2007) using Kyoto Encyclopedia of Genes and Genomes (KEGG) and GO gene sets from MSigDB.

#### Quantitative proteomics

Stable isotope labeling by amino acids in cell culture (SILAC) was performed using light (^12^C_6_-Arg/^12^C_6_-Lys) or heavy (^13^C_6_-Arg/^13^C_6_-Lys) arginine and lysine (Ong et al. 2002; Colaert et al. 2010). Cells were fully labeled over 2 weeks in SILAC medium, with *NBPF1* expression induced by Dox in heavy-labeled cells. Equal amounts of light- and heavy-labeled protein lysates were mixed, digested with trypsin, fractionated by HPLC, and analyzed by liquid chromatography-tandem mass spectrometry (LC-MS/MS) on an LTQ Orbitrap instrument. Protein ratios were quantified and subjected to GO enrichment analysis.

## Results

### Transcriptome analysis

RNA-seq analysis of doxycycline-induced *NBPF1*-overexpressing DLD1Tr21 cells revealed strong, specific downregulation of mitochondrial genes relative to uninduced controls (Fig. 1; Table 1; Tables S1-S3). Top enriched GO terms among downregulated genes included mitochondrial inner membrane, mitochondrion, and respiratory chain components (all FDR < 0.05). No significant mitochondrial enrichment was observed in doxycycline-treated mock control cells. Notably, the sample with the highest *NBPF1* expression (as determined by EGFP levels) showed the strongest downregulation, consistent with a dosage-dependent effect (Fig S2; Table S4).

**Table 1.**
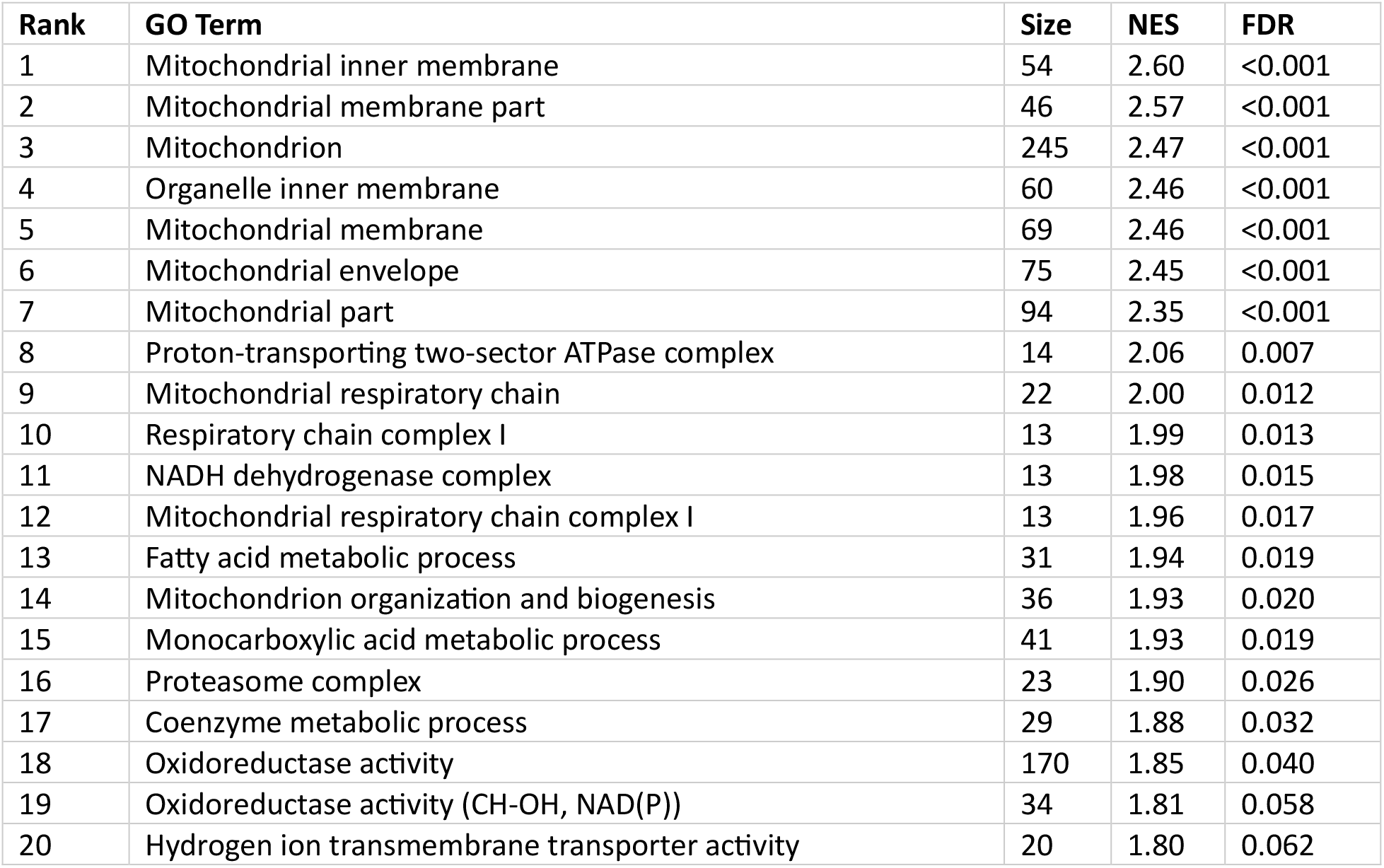
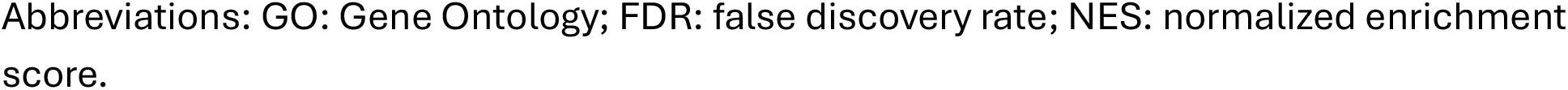
Top 20 downregulated GO terms in DLD1Tr21/NBPF cells (RNA-seq).

| Rank | GO Term | Size | NES | FDR |
| --- | --- | --- | --- | --- |
| 1 | Mitochondrial inner membrane | 54 | 2.60 | <0.001 |
| 2 | Mitochondrial membrane part | 46 | 2.57 | <0.001 |
| 3 | Mitochondrion | 245 | 2.47 | <0.001 |
| 4 | Organelle inner membrane | 60 | 2.46 | <0.001 |
| 5 | Mitochondrial membrane | 69 | 2.46 | <0.001 |
| 6 | Mitochondrial envelope | 75 | 2.45 | <0.001 |
| 7 | Mitochondrial part | 94 | 2.35 | <0.001 |
| 8 | Proton-transporting two-sector ATPase complex | 14 | 2.06 | 0.007 |
| 9 | Mitochondrial respiratory chain | 22 | 2.00 | 0.012 |
| 10 | Respiratory chain complex I | 13 | 1.99 | 0.013 |
| 11 | NADH dehydrogenase complex | 13 | 1.98 | 0.015 |
| 12 | Mitochondrial respiratory chain complex I | 13 | 1.96 | 0.017 |
| 13 | Fatty acid metabolic process | 31 | 1.94 | 0.019 |
| 14 | Mitochondrion organization and biogenesis | 36 | 1.93 | 0.020 |
| 15 | Monocarboxylic acid metabolic process | 41 | 1.93 | 0.019 |
| 16 | Proteasome complex | 23 | 1.90 | 0.026 |
| 17 | Coenzyme metabolic process | 29 | 1.88 | 0.032 |
| 18 | Oxidoreductase activity | 170 | 1.85 | 0.040 |
| 19 | Oxidoreductase activity (CH-OH, NAD(P)) | 34 | 1.81 | 0.058 |
| 20 | Hydrogen ion transmembrane transporter activity | 20 | 1.80 | 0.062 |

Abbreviations: GO: Gene Ontology; FDR: false discovery rate; NES: normalized enrichment score.

**Fig. 1.**
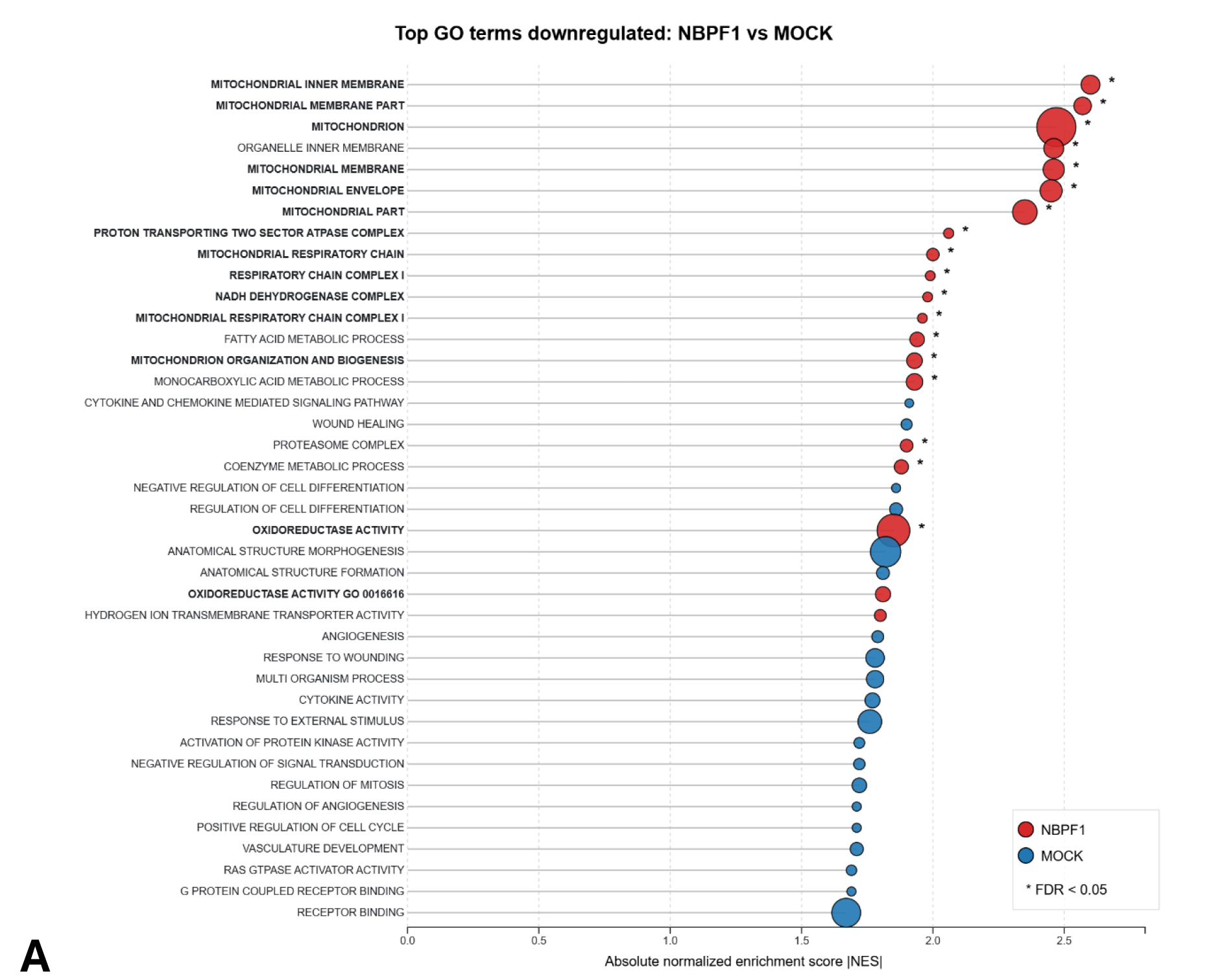
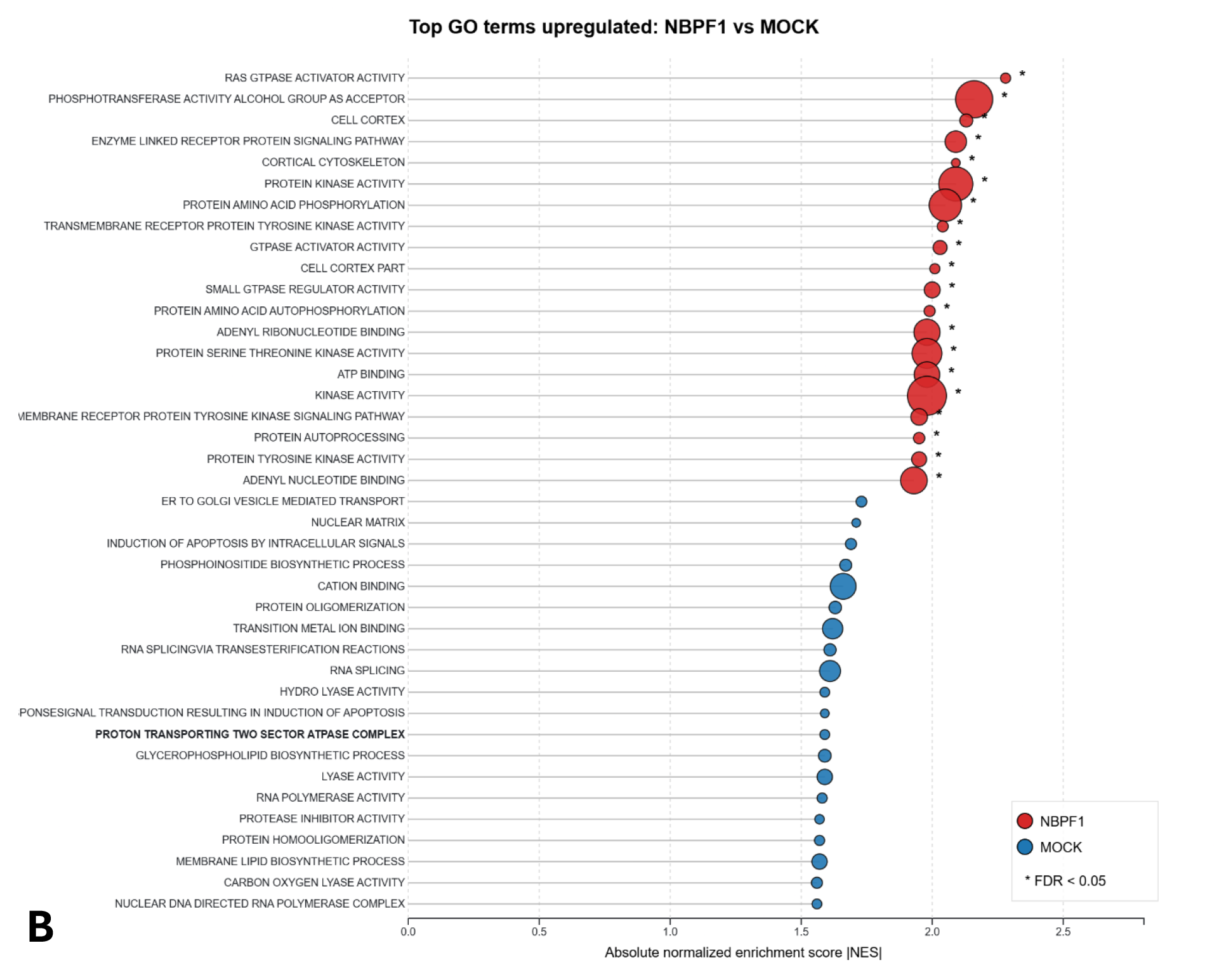
Lollipop plots showing the top 20 downregulated (A) and upregulated (B) GO terms in DLD1Tr21/NBPF versus DLD1Tr21/Mock cells (RNA-seq GSEA analysis). Normalized enrichment scores (NES) for the top 20 downregulated Gene Ontology (GO) terms in each condition were combined and ranked by NES. Each dot represents one GO term. X-axis: NES, with higher values indicating stronger downregulation (plotted as absolute values for visual comparison between plots). Y-axis: GO biological process/cellular component terms (ranked by descending NES). Dot size: Proportional to the number of genes in the gene set (larger dots represent larger gene sets). Color: Red dots = DLD1Tr21/NBPF dataset; blue dots = DLD1Tr21/Mock dataset. Terms related to mitochondrial function are in boldface.

### Proteomic analysis

SILAC-based quantitative mass spectrometry identified 79 differentially expressed proteins upon *NBPF1* induction (99% confidence), of which 52 were downregulated (Tables S5-S6). GO enrichment analysis showed highly significant overrepresentation of mitochondrial electron transport chain components (adjusted p = 1.81 × 10^−11^) and NADH dehydrogenase activity (adjusted p = 2.43 × 10^−11^) among downregulated proteins. Importantly, eight of the top 24 downregulated proteins (including UQCRC1, PHB, PHB2, IMMT, NDUFS2, NDUFS8, NDUFS1, and UQCRB) overlapped with the RNA-seq dataset and have well-established roles in mitochondrial structure and function.

### Live-cell imaging

To validate these findings functionally, we performed live-cell imaging using MitoTracker CMXRos. *NBPF1*-overexpressing cells displayed markedly reduced mitochondrial signal, with significant decreases in red fluorescence coverage, intensity, and object count compared to uninduced or mock controls (Fig. 2; Figs S3-S5). This effect was clearly cell-density dependent—minimal at low density but pronounced near confluency (Fig S6). Treatment with doxycycline alone had no effect on mitochondrial parameters in mock cells, confirming specificity to *NBPF1*.

**Fig. 2.**
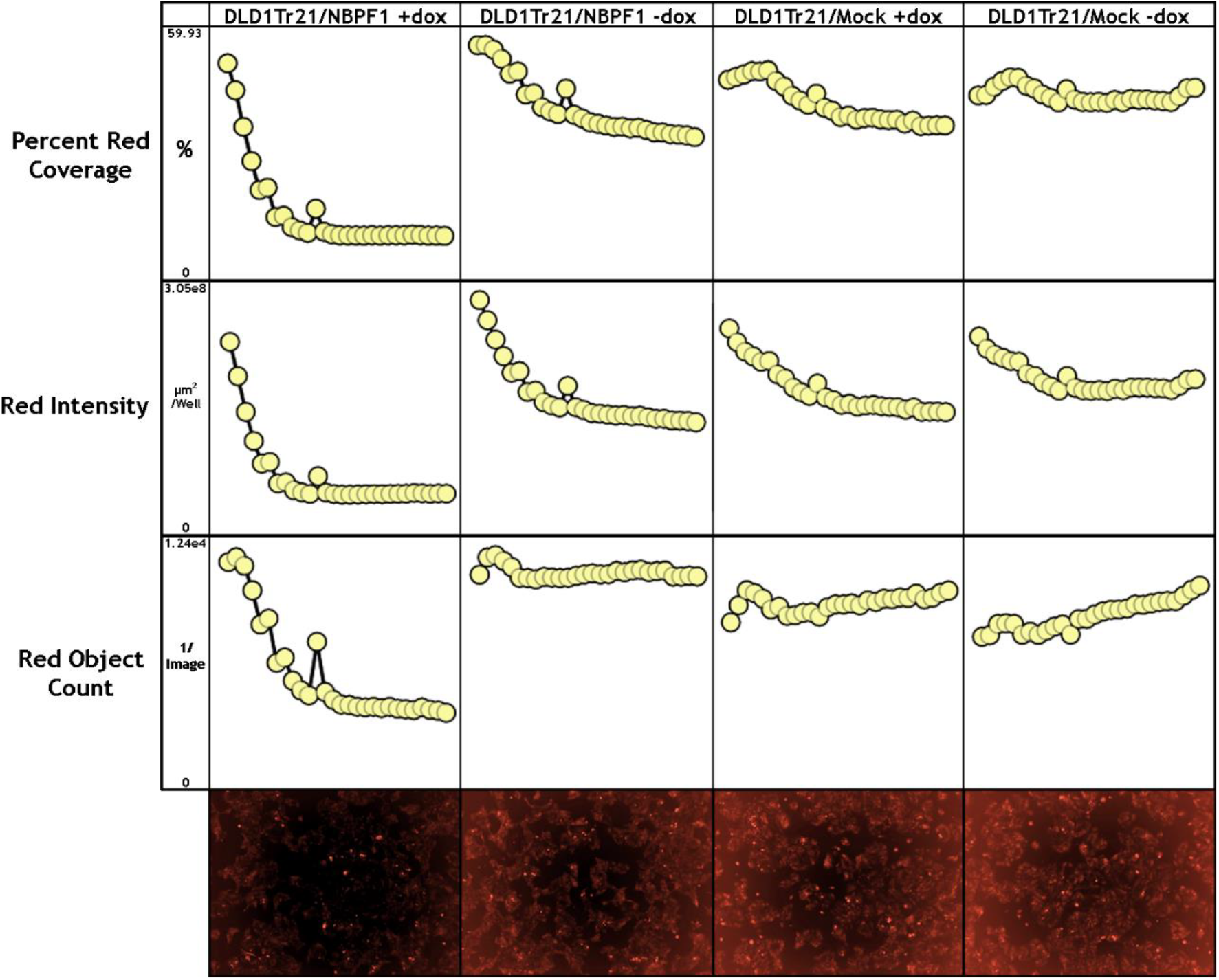
DLD1-NBPF1 or DLD1-MOCK cells incubated with MitoTracker, a reagent that binds to mitochondria and fluoresces red. Each circle represents a time point at 4 h. Red coverage (confluency), average red object intensity, and red object count are plotted against time over the course of 4 days from the time of doxycycline application for each cell line with and without doxycycline to initiate transcription of the transgene. A low magnification example image from each condition is shown below the plots to indicate general levels of red signal.

## Discussion

This study provides the first direct evidence that Olduvai domains downregulate mitochondrial pathways. Transcriptomic, proteomic, and live-cell imaging data demonstrate that *NBPF1* overexpression reduces electron transport chain activity, mitochondrial abundance, and related functions. The effect is dosage sensitive and cell-density dependent, becoming strongest near confluency.

### Density-dependent control of neuronal size and scaling

By lowering cellular metabolic state, Olduvai domains would be expected to favor prolonged progenitor proliferation at the expense of rapid differentiation and growth. Because cell growth is highly energy demanding, reduced mitochondrial activity could limit cell growth and thereby help maintain smaller, more uniform neuronal size. This shift offers a plausible explanation for the distinctive primate scaling rule of near-constant neuronal density despite increasing brain size (Herculano-Houzel 2009), in contrast to most mammals, where brain enlargement is accompanied by less densely packed (and therefore presumably larger) neurons.

Recent work strengthens this interpretation. Energy depletion and a reduced ATP/ADP ratio induce expression of Olduvai-containing NBPF proteins (Ekvik et al. 2026). Importantly, we find that Olduvai-mediated mitochondrial damping is density dependent, occurring primarily at high cell densities where local energy stress is greatest. Together, these findings support a model in which Olduvai acts as a tunable metabolic brake: high cell density triggers energy stress that induces Olduvai expression, which in turn attenuates mitochondrial activity.

This feedback loop may slow developmental tempo, extend the period of neurogenesis, and restrict cell growth—potentially enabling the generation of more neurons while preserving a low-energy state and relatively uniform neuronal size, a hallmark of primate cortical architecture.

### Brain evolution and neoteny

Neoteny is a delay in the timing of developmental events in one species relative to another, resulting in the prolonged retention of juvenile or infantile characteristics into adulthood. It has long been considered a major evolutionary mechanism in primate and human evolution (Gould 1977; Thiessen 1997). A growing body of work indicates that mitochondrial metabolism can act as a developmental pacemaker, with reduced mitochondrial activity slowing cellular maturation and promoting neoteny (Namba et al. 2020; Iwata et al. 2023; Casimir et al. 2024; Lindhout et al. 2024; Liu and Auwerx 2023). Our proteomics data show that Olduvai expression most strongly downregulates electron transport chain and NADH-related mitochondrial proteins—the same pathways implicated in controlling species-specific developmental timing (Diaz-Cuadros et al. 2023).

Given these findings, we suggest that the Olduvai-mediated suppression of mitochondrial energy production may therefore slow cellular maturation and extend developmental windows. In the brain, this could prolong the neurogenic phase, enabling increased neuron production. More broadly, because Olduvai domains are expressed across multiple tissues, this mechanism may contribute to generalized neoteny rather than acting solely on brain expansion. Consistent with this idea, Olduvai copy number increases progressively across primate lineages in parallel with both brain size and neoteny (Fig. 3 and Table S7; O’Bleness et al. 2012; Gould 1977; Thiessen 1997; Somel et al. 2009). In addition, Olduvai-minus (knockout [KO]) mice that lack their single ancestral Olduvai copy exhibit hyperactivity (Keeney et al. 2015b), as might be expected from reduced metabolic restraint.

**Fig. 3.**
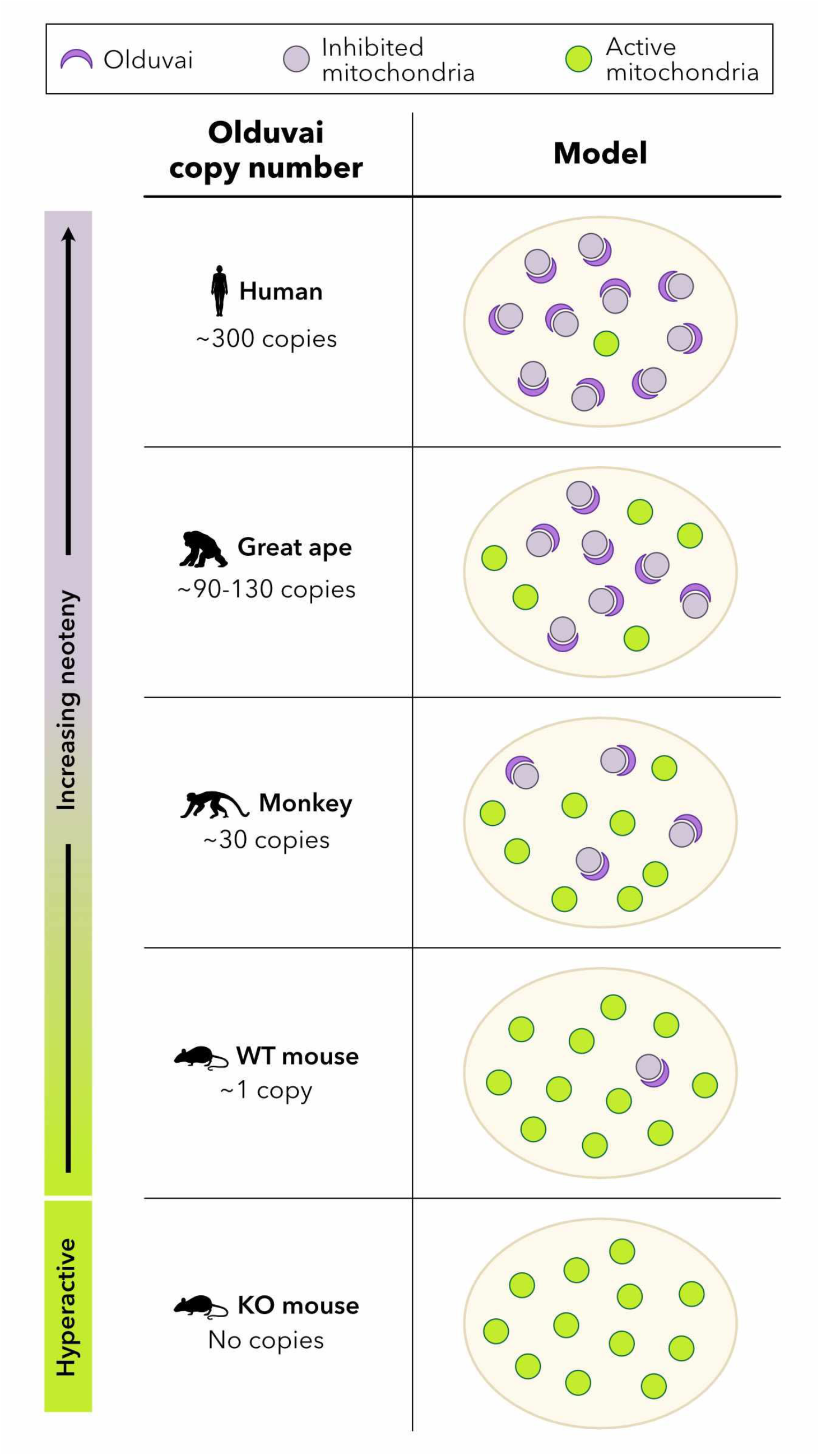
Model linking Olduvai copy number, mitochondrial inhibition, and neoteny. The schematic illustrates a hypothesized mechanism in which Olduvai proteins inhibit mitochondrial function in a copy-number-dependent manner. This inhibition is proposed to slow developmental timing, thereby promoting neoteny across species in proportion to Olduvai copy number (Gould 1977; Thiessen 1997). The hyperactivity phenotype observed in Olduvai knockout (KO) mice is also indicated (Keeney, et al. 2015).

### Pairing of Olduvai and *NOTCH2NL*

The Olduvai findings reported here also have potential relevance to human-specific *NOTCH2NL* genes, which promote cortical neurogenesis (Fiddes et al. 2018; Suzuki et al. 2018; Florio et al. 2018). The majority of human-specific Olduvai copies are encoded by three *NBPF* genes that lie adjacent to, and are tightly co-regulated with, three *NOTCH2NL* genes (Fiddes et al. 2019). These *NBPF*/*NOTCH2NL* pairs evolved jointly as two-gene units and have undergone three successive duplications that recently added 119 Olduvai copies to the human genome. While *NOTCH2NL* paralogs enhance Notch signaling and promote apical progenitor proliferation, their paired *NBPF* genes have been shown to drive basal radial glia expansion (Eşiyok et al., 2025), a population of developing neurons whose abundance and proliferative capacity are greatly expanded in primates. Basal radial glia are intriguing in that they’ve lost ventricular attachment, yet they still behave like progenitor cells.

In this context, Olduvai-mediated mitochondrial downregulation may act as an energy-sensitive modulator that slows maturation and helps prevent premature differentiation in basal progenitors. Acting in a complementary manner, these paired modules—*NOTCH2NL* promotes proliferation while Olduvai restrains differentiation—may prolong the progenitor pool and support human-specific cortical expansion. Such a coordinated, synergistic mechanism could explain why these Olduvai–*NOTCH2NL* pairs have been strongly selected for in recent human evolution.

## Conclusions

In summary, these data suggest that Olduvai domains function as tunable, dosage-dependent regulators linking cellular energetic state to developmental timing. Through copy-number expansion and metabolic restraint, they appear to promote both general neoteny and extended neurogenesis, thereby increasing neuron number and contributing to the evolutionary expansion of brain size that characterizes the human lineage. Identifying the precise molecular intermediaries connecting Olduvai domains to mitochondrial regulation remains an important next step.

## Supporting information

Keeney_Sikela_Supp files

## Statements and Declarations

## Acknowledgments

We thank Mark Johnston, Chet Sherwood, Santiago Fregoso and Jeff Moore for helpful discussions. We also thank Julie Ho Sung and Natasia Paukovich for help with illlustrations and Julie Fiorilla for help with manuscript preparation.

## Author Contributions

J.K. and J.M.S. conceived the project. J.K. performed the majority of the experiments, and the key findings of this study form part of his PhD thesis (2014). J.M.S. supervised the project. Material preparation, data collection, and analysis were performed by:

- RNA-seq: D.A. and K.J.
- Proteomics: A.S., K.G., and J.V.
- Cell line generation and culture: V.A. and K.V.
- Live-cell imaging and mitochondrial measurements: J.K., P.L. and M.P.

## Additional contributions

- V.A., K.V., N.A., J.D., P.L., S.F., N.P,, B.V., and F.V. provided critical insights into experimental strategies, data interpretation, and project feedback.

J.K. and J.M.S. wrote the original draft of the manuscript. All authors commented on previous versions of the manuscript and read and approved the final manuscript.

