## Supplementary material for "Olduvai Domains Downregulate Mitochondrial Pathways to Promote Human Brain Evolution and Neoteny": Keeney_Sikela_Supp files

Update Note

This preprint is a revised and updated version of the manuscript previously posted on Research Square. It replaces the earlier version. The previous version remains available here:

https://doi.org/10.21203/rs.3.rs-10345985/v1

DOI: 10.21203/rs.3.rs-10345985/v1

This manuscript is currently under review at *Human Genetics*.

This is a revision of our 2024 bioRxiv preprint:

https://doi.org/10.1101/2024.10.21.619278

### Olduvai Domains Downregulate Mitochondrial Pathways to Promote Human Brain Evolution and Neoteny

Jonathon G. Keeney,[1,*] David Astling,[1] Vanessa Andries,[2,3,#] Karl Vandepoele,[2,3,$] Nathan Anderson,[1] Jonathan M. Davis,[1] Pamela Lopert,[4] Manisha Patel,[4] Kevin Jones,[1] Jonathan Vandenbussche,[5,6,&] Kris Gevaert,[5,6] An Staes,[5,6,7] Santos Franco,[8] Natasia Paukovich,[1] Beat Vögeli,[1] Frans van Roy,[2,3] and James M. Sikela[1,9,†,^]

[1]Department of Biochemistry and Molecular Genetics, University of Colorado Anschutz Medical Campus, Aurora, CO 80045, USA
[2]Inflammation Research Center, VIB, 9052 Ghent, Belgium
[3]Department of Biomedical Molecular Biology, Ghent University, 9052 Ghent, Belgium
[4]Department of Pharmaceutical Sciences, School of Pharmacy, University of Colorado Anschutz Medical Campus, Aurora, CO 80045, USA
[5]Department of Biomolecular Medicine, Ghent University, 9052 Ghent, Belgium
[6]VIB Center for Medical Biotechnology, VIB, 9052 Ghent, Belgium
[7]VIB Proteomics Core, 9052 Ghent, Belgium
[8]Department of Pediatrics, University of Colorado Anschutz Medical Campus, Aurora, CO 80045, USA
[9]Human Medical Genetics and Neuroscience Programs, University of Colorado Anschutz Medical Campus, Aurora, CO 80045, USA
*Current address: Department of Biochemistry and Molecular Medicine, George Washington University of Medicine and Health Sciences, Washington, DC 20037, USA
[#]Current address: Department of Internal Medicine and Pediatrics, Ghent University, Ghent, Belgium
[$]Current address: Department of Clinical Chemistry, Ghent University Hospital, 9000 Ghent, Belgium
[&]Current address: RIC group, P. Kennedypark 6, 8500 Kortrijk, Belgium
† A brief Forum article by the corresponding author on aspects of this topic is in press at *Trends in Genetics*.

#### Supplementary Methods

##### Cell culture and live-cell imaging

Doxycyclin hydrochloride (Dox; Sigma, Grand Island, New York) was used at a final concentration of 2 µg/ml to induce expression of either NBPF1-IRES-EGFP or EGFP in DLD1Tr21/NBPF1 and DLD1Tr21/Mock cells, respectively. DLD1Tr21/Mock and DLD1Tr21/NBPF1 cells were described previously (Vandepoele et al., 2008). Briefly, in the colon cancer cell line DLD1Tr21 Tet-On originally obtained from Van de Wetering et al., 2002, a Tet-On system activates transcription of the gene of interest in the presence of Dox. The cDNA for NBPF1-IRES-EGFP, fused to an amino-terminal flag tag, was cloned in the pcDNA4/TO vector (Invitrogen). This construct was stably transfected in DLD1Tr21 cells and subcloned after fluorescence-activated cell sorting. A final clone was more than 90% positive for flag-tagged *NBPF1* expression upon induction with Dox for 48 h. For mitochondrial analysis, media was removed at 24 h after plating and replaced with media including 200 nM MitoTracker CMSROX (Gibco) and either 2 µg/ml Dox or no Dox. Cells were then cultured at 37$^{o}$C in 5% $CO_2$ in an IncuCyte kinetic life cell imaging system (Essen Bioscience, Ann Arbor, MI). Red (MitoTracker), green (from IRES-EGFP on the construct, indicating successful Dox induction), and phase contrast images were collected every 4 h.

##### Library preparation and Illumina sequencing

For both DLD1Tr21/NBPF1 and DLD1Tr21/Mock cells, eight T-175 flasks (Falcon) were cultured at 37$^{o}$C in 5% CO2. After 24 h, media was changed. Four flasks of each cell line were refreshed with medium containing Dox, whereas the medium of the remaining four flasks was refreshed with medium only. After 24 h, cells were trypsinized (Gibco) and collected in complete medium. cDNA libraries were constructed starting from 1 µg total RNA. PolyA-plus mRNAs were selected for by using oligo dT beads according to the manufacturer's instructions, then reverse transcribed to cDNA and prepared for next-generation sequencing according to the manufacturer's protocol (Illumina TruSeq RNA Sample Preparation v2 Guide). The cDNA libraries were validated on the Agilent 2100 Bioanalyzer using DNA-1000 chip. Cluster generation was done on the Illumina cBot using a Single Read Flow Cell with a Single Read cBot reagent plate (TruSeq SR Cluster Kit). Sequencing of the clustered flow cell was performed on the Illumina HiSeq 2000 using TruSeq SBS v3 reagents. The sequencer was programmed with a single read at 100 cycles. Sequencing images were generated through the sequencing platform (Illumina HiSeq, 2000). The raw data were analyzed in four steps: image analysis, base calling, sequence alignment, and variant analysis and counting. An additional step was required to convert the base call files (.bcl) into *_qseq.txt files. For multiplexed lanes/samples, a de-multiplexing step was performed before the alignment step.

**RNA-seq data analysis**

On average, we obtained 20 million (range = 12–30 million) single-end 100 bp sequencing reads per sample. Reads were mapped against the human genome using Tophat (version 2.0.9) (Kim et al., 2013) and the NCBI reference annotation (build 37.2) as a guide. We allowed 3 mismatches for the initial alignment and 2 mismatches per segment with 25-bp segments. On average, 90% (85%–94%) of the reads aligned to the human genome. Next, we employed Cufflinks (version 2.1.1) (Trapnell et al., 2010) to assemble the transcripts using the RefSeq annotations as the guide but allowing for novel isoform discovery in each sample. Isoforms were ignored if the number of supporting reads was less than 30 and if the isoform fraction was less than 10% for the gene. The data were fragment bias corrected, multi-read corrected, and normalized by the total number of reads. The transcript assemblies for each sample were merged using Cuffmerge (Trapnell et al., 2010). We next computed the transcripts' FPKM values (Fragments Per Kilobase of exon per Million fragments mapped) by rerunning Cufflinks using the merged assembly as the guide. Gene expression was estimated by summing the FPKM values of multiple transcripts that represent the same gene. All other analyses were performed in R/Bioconductor (R version 3.0.1; R Development Core, 2011).

Significantly changed genes were identified using the R package LIMMA (Smyth, 2005). Significant genes were identified using the lmFit, eBayes, and decideTests functions (FDR < 0.05 and fold change > 1.5) from the LIMMA package. The statistics were moderated by LIMMA using empirical Bayes shrinkage (via the eBayes function), global multiple testing strategy, and Benjamini & Hochberg adjustment. Gene set enrichment analysis (GSEA) was conducted using the Kyoto Encyclopedia of Genes and Genomes (KEGG) and Gene Ontology (GO) gene set definitions obtained from the Molecular Signatures Database v.3.1. Enriched pathways were identified by running GSEA using 1,000 permutations as a standalone Java app (version 2.0.13; Subramanian et al., 2007).

**Proteomics**

Stable isotope labeling by amino acids in cell culture (SILAC) with heavy arginine and lysine was used for mass spectrometry-based quantitative proteomic analysis (Colaert et al., 2010). However, its accuracy is compromised by the metabolic conversion of arginine to proline in eukaryotes. As a result, $^{13}C_6$-Arg gets partly converted into $^{13}C_5$-Pro, which results in the generation of multiple satellite peaks for Pro-containing tryptic peptides in the labeled state. This hampers accurate quantitation. We therefore optimized the $^{13}C_6$-Arg concentration of the DLD1Tr21 medium and determined that 30 mg $^{13}C_6$-Arg per l medium was optimal to minimize $^{13}C_6$-Arg conversion and that the cells had to be grown for 2 weeks in SILAC medium to obtain full labeling of the cells.

Medium for light labeled cells (− Dox) was RPMI-1640 medium (without Arg and Lys), supplemented with GlutaMax (Gibco), 10% dialyzed fetal calf serum, 0.03 mg/ml $^{12}C_6$-Arg and 0.03 mg/ml $^{12}C_6$-Lys. Medium for heavy labeled cells (+ Dox) was RPMI-1640 medium (without Arg and Lys), supplemented with GlutaMax, 10% dialyzed fetal calf serum, 0.03 mg/ml $^{13}C_6$-Arg and 0.03 mg/ml $^{13}C_6$-Lys. Dox for the induction of *NBPF1* expression was added to the medium at a final concentration of 2 µg/ml.

Cells were grown for 2 weeks in the appropriate SILAC medium, and cell pellets were then resuspended in 0.5 ml of cell lysis buffer containing 50 mM sodium phosphate pH 7.5, 100 mM NaCl, 0.8% CHAPS (w/v), and Complete Protease Inhibitor cocktail (Roche). Cells were lysed on ice for 30 min, and the insoluble fraction was removed by centrifugation (15 min at 16,000 g at 4°C). Protein measurements of the supernatant occurred by use of the DC Protein Assay kit (Bio-Rad).

The light and heavy labeled samples were mixed in a 1:1 ratio (1.7 mg of protein each). 380 mg guanidinium hydrochloride was added, and the sample volume was adjusted to 1 ml by adding lysis buffer to obtain a final guanidinium hydrochloride concentration of about 4 M.

S-reduction and S-alkylation were performed by adding 26 µl of freshly prepared and pH-adjusted 570-mM TCEP.HCl solution and 40 µl of freshly prepared 750 mM iodoacetamide solution, obtaining a final concentration of 15 mM TCEP.HCl and 30 mM iodoacetamide. The reaction proceeded for 15 min while mixing at 30°C in the dark. The sample was then desalted on a NAP-10 column in 20 mM ammonium bicarbonate pH 7.6. The protein concentration in the NAP-10 eluate was measured using the Bio-Rad DC Protein Assay Kit, and 750 µl (0.868 mg protein) was taken to carry out the next steps. The sample was boiled for 5 min and directly transferred to ice for 10 min. Trypsin was added in a trypsin/protein ratio of 1/50 (w/w), and the sample was incubated overnight at 37°C.

The peptide mixture was dried in a Speedvac and then redissolved in 1% acetic acid. After checking the pH, the sample was loaded on an HPLC column and 1 min fractions were collected between 20 and 80 min. Fractions with a retention time difference of 20 min were pooled, dried down, and analyzed with the LTQ-Orbitrap mass spectrometer. GO analysis was performed with WebGestalt V2 (Zhang et al., 2005) comparing the obtained results to the complete human proteome. The distribution of ratio values (light/heavy) was analyzed using a base 2 logarithmic scale, which is a common practice for protein expression data (MacCoss et al., 2003).

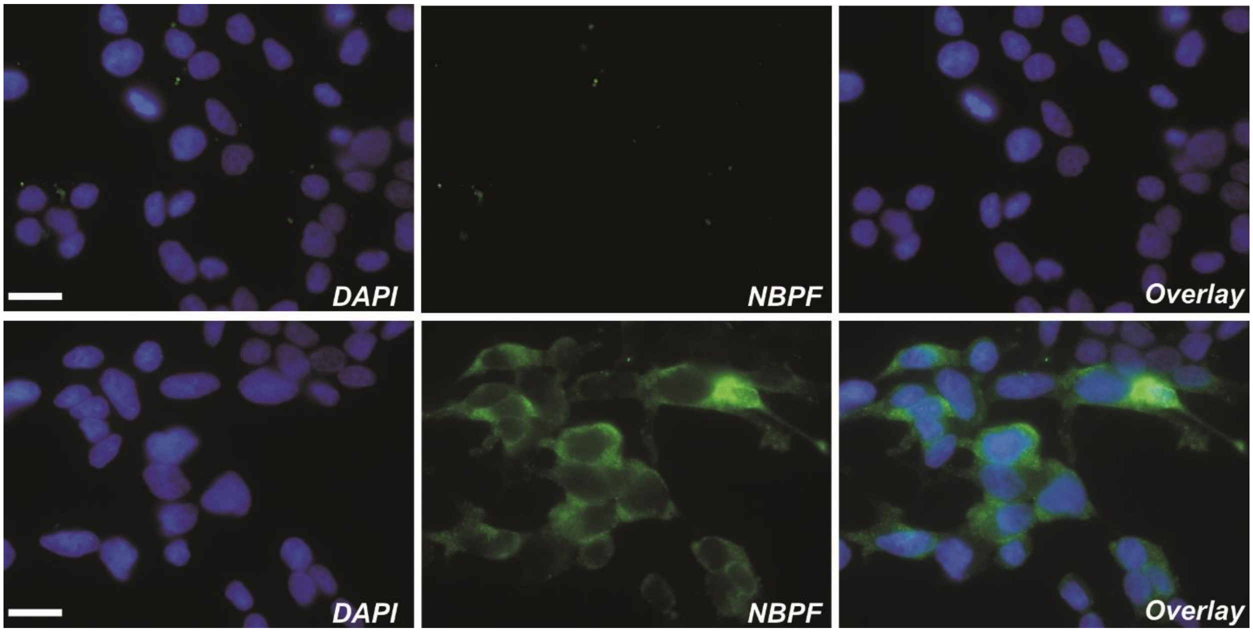


**Supplemental Figure 1. Immunofluorescent analysis of DLD1Tr21/NBPF1 cell cultures, used for proteomic analysis**

DLD1Tr21/NBPF1 cells in the absence of dox (top panels) were grown for 2 weeks in medium with light $^{12}C_6$-Lys/Arg amino acids, whereas DLD1Tr21/NBPF1 cells in the presence of dox (lower panels) were grown in medium with heavy $^{13}C_6$-Lys/Arg amino acids. Immunofluorescent staining with an anti-NBPF antibody (sc-88241) showed that *NBPF1* was only expressed in cells that were grown in medium with heavy $^{13}C_6$-Lys/Arg amino acid precursors in the presence of dox (lower panels). Nuclei were visualized with DAPI. Scale bars, 25 µm.

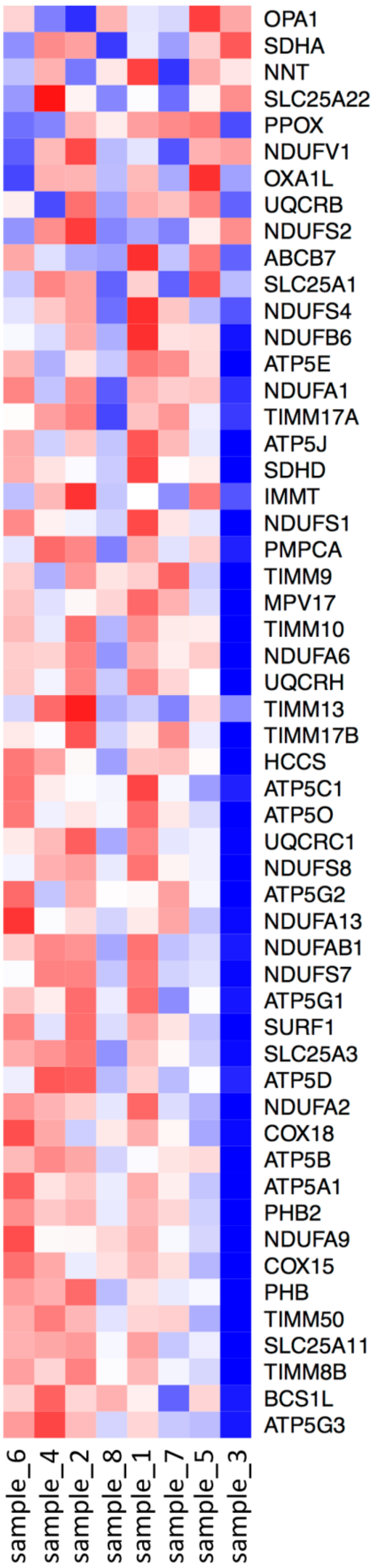

OPA1
SDHA
NNT
SLC25A22
PPOX
NDUFV1
OXA1L
UQCRB
NDUFS2
ABCB7
SLC25A1
NDUFS4
NDUFB6
ATP5E
NDUFA1
TIMM17A
ATP5J
SDHD
IMMT
NDUFS1
PMPCA
TIMM9
MPV17
TIMM10
NDUFA6
UQCRH
TIMM13
TIMM17B
HCCS
ATP5C1
ATP5O
UQCRC1
NDUFS8
ATP5G2
NDUFA13
NDUFAB1
NDUFS7
ATP5G1
SURF1
SLC25A3
ATP5D
NDUFA2
COX18
ATP5B
ATP5A1
PHB2
NDUFA9
COX15
PHB
TIMM50
SLC25A11
TIMM8B
BCS1L
ATP5G3
sample_6
sample_4
sample_2
sample_8
sample_1
sample_7
sample_5
sample_3

**Supplemental Figure 2. Heatmap showing the transcript level changes detected in RNA-seq**

Four replicates each of DLD1Tr21/NBPF1 (odd numbered samples) and DLD1Tr21/Mock (even numbered samples) are listed at the bottom, and detected transcripts with the strongest significance are listed to the right. Red indicates an increased detection and blue indicates a reduced detection, with stronger values indicated by a more intense color. Values are obtained by comparing each induced replicate (+ doxycycline) with each uninduced replicate (− doxycycline).

#### Top 20 GO terms downregulated in DLD1Tr21/Mock upon RNA-seq analysis

| | Pathway | Size | NES | FDR |
|---|---|---|---|---|
| 1 | CYTOKINE_AND_CHEMOKINE_MEDIATED_SIGNALING_ PATHWAY | 10 | 1.91 | 0.259 |
| 2 | WOUND_HEALING | 17 | 1.90 | 0.152 |
| 3 | NEGATIVE_REGULATION_OF_CELL_DIFFERENTIATION | 11 | 1.86 | 0.167 |
| 4 | REGULATION_OF_CELL_DIFFERENTIATION | 24 | 1.86 | 0.134 |
| 5 | ANATOMICAL_STRUCTURE_MORPHOGENESIS | 146 | 1.82 | 0.189 |
| 6 | ANATOMICAL_STRUCTURE_FORMATION | 24 | 1.81 | 0.173 |
| 7 | ANGIOGENESIS | 20 | 1.79 | 0.189 |
| 8 | RESPONSE_TO_WOUNDING | 52 | 1.78 | 0.172 |
| 9 | MULTI_ORGANISM_PROCESS | 45 | 1.78 | 0.164 |
| 10 | CYTOKINE_ACTIVITY | 33 | 1.77 | 0.162 |
| 11 | RESPONSE_TO_EXTERNAL_STIMULUS | 87 | 1.76 | 0.157 |
| 12 | ACTIVATION_OF_PROTEIN_KINASE_ACTIVITY | 16 | 1.72 | 0.220 |
| 13 | NEGATIVE_REGULATION_OF_SIGNAL_TRANSDUCTION | 18 | 1.72 | 0.216 |
| 14 | REGULATION_OF_MITOSIS | 31 | 1.72 | 0.204 |
| 15 | REGULATION_OF_ANGIOGENESIS | 11 | 1.71 | 0.207 |
| 16 | POSITIVE_REGULATION_OF_CELL_CYCLE | 11 | 1.71 | 0.200 |
| 17 | VASCULATURE_DEVELOPMENT | 25 | 1.71 | 0.188 |
| 18 | RAS_GTPASE_ACTIVATOR_ACTIVITY | 15 | 1.69 | 0.202 |
| 19 | G_PROTEIN_COUPLED_RECEPTOR_BINDING | 11 | 1.69 | 0.205 |
| 20 | RECEPTOR_BINDING | 131 | 1.67 | 0.222 |

**Supplemental Table 1. Pathways downregulated based on RNA-seq analysis of the mock cell line after treatment with doxycycline**

#### Top 20 GO terms upregulated in DLD1Tr21/NBPF upon RNA-seq analysis

| | Pathway | Size | NES | FDR |
|---|---|---|---|---|
| 1 | RAS_GTPASE_ACTIVATOR_ACTIVITY | 15 | -2.28 | 0.004 |
| 2 | PHOSPHOTRANSFERASE_ACTIVITY_ALCOHOL_GROUP_AS_ACCEPTOR | 208 | -2.16 | 0.018 |
| 3 | CELL_CORTEX | 25 | -2.13 | 0.025 |
| 4 | ENZYME_LINKED_RECEPTOR_PROTEIN_SIGNALING_PATHWAY | 69 | -2.09 | 0.032 |
| 5 | CORTICAL_CYTOSKELETON | 11 | -2.09 | 0.026 |
| 6 | PROTEIN_KINASE_ACTIVITY | 175 | -2.09 | 0.022 |
| 7 | PROTEIN_AMINO_ACID_PHOSPHORYLATION | 157 | -2.05 | 0.032 |
| 8 | TRANSMEMBRANE_RECEPTOR_PROTEIN_TYROSINE_KINASE_ACTIVITY | 18 | -2.04 | 0.029 |
| 9 | GTPASE_ACTIVATOR_ACTIVITY | 29 | -2.03 | 0.029 |
| 10 | CELL_CORTEX_PART | 15 | -2.01 | 0.035 |
| 11 | SMALL_GTPASE_REGULATOR_ACTIVITY | 38 | -2.00 | 0.037 |
| 12 | PROTEIN_AMINO_ACID_AUTOPHOSPHORYLATION | 18 | -1.99 | 0.038 |
| 13 | ADENYL_RIBONUCLEOTIDE_BINDING | 103 | -1.98 | 0.040 |
| 14 | PROTEIN_SERINE_THREONINE_KINASE_ACTIVITY | 134 | -1.98 | 0.039 |
| 15 | ATP_BINDING | 98 | -1.98 | 0.037 |
| 16 | KINASE_ACTIVITY | 229 | -1.98 | 0.035 |
| 17 | TRANSMEMBRANE_RECEPTOR_PROTEIN_TYROSINE_KINASE_SIGNALING_PATHWAY | 41 | -1.95 | 0.044 |
| 18 | PROTEIN_AUTOPROCESSING | 19 | -1.95 | 0.042 |
| 19 | PROTEIN_TYROSINE_KINASE_ACTIVITY | 33 | -1.95 | 0.040 |
| 20 | ADENYL_NUCLEOTIDE_BINDING | 107 | -1.93 | 0.043 |

**Supplemental Table 2. Pathways found to be upregulated in our RNA-seq analysis after exogenous expression of *NBPF1***

#### Top 20 GO terms upregulated in DLD1Tr21/Mock upon RNA-seq analysis

| | Pathway | Size | NES | FDR |
|---|---|---|---|---|
| 1 | ER_TO_GOLGI_VESICLE_MEDIATED_TRANSPORT | 17 | -1.73 | 1.000 |
| 2 | NUCLEAR_MATRIX | 11 | -1.71 | 1.000 |
| 3 | INDUCTION_OF_APOPTOSIS_BY_INTRACELLULAR_SIGNALS | 18 | -1.69 | 1.000 |
| 4 | PHOSPHOINOSITIDE_BIOSYNTHETIC_PROCESS | 21 | -1.67 | 1.000 |
| 5 | CATION_BINDING | 100 | -1.66 | 0.923 |
| 6 | PROTEIN_OLIGOMERIZATION | 23 | -1.63 | 0.986 |
| 7 | TRANSITION_METAL_ION_BINDING | 62 | -1.62 | 0.900 |
| 8 | RNA_SPLICINGVIA_TRANSESTERIFICATION_REACTIONS | 22 | -1.61 | 0.836 |
| 9 | RNA_SPLICING | 66 | -1.61 | 0.752 |
| 10 | HYDRO_LYASE_ACTIVITY | 14 | -1.59 | 0.754 |
| 11 | DNA_DAMAGE_RESPONSESIGNAL_TRANSDUCTION_RESULTING_IN_INDUCTION_OF_APOPTOSIS | 11 | -1.59 | 0.701 |
| 12 | PROTON_TRANSPORTING_TWO_SECTOR_ATPASE_COMPLEX | 14 | -1.59 | 0.645 |
| 13 | GLYCEROPHOSPHOLIPID_BIOSYNTHETIC_PROCESS | 24 | -1.59 | 0.597 |
| 14 | LYASE_ACTIVITY | 35 | -1.59 | 0.558 |
| 15 | RNA_POLYMERASE_ACTIVITY | 15 | -1.58 | 0.553 |
| 16 | PROTEASE_INHIBITOR_ACTIVITY | 13 | -1.57 | 0.563 |
| 17 | PROTEIN_HOMOOLIGOMERIZATION | 15 | -1.57 | 0.546 |
| 18 | MEMBRANE_LIPID_BIOSYNTHETIC_PROCESS | 35 | -1.57 | 0.517 |
| 19 | CARBON_OXYGEN_LYASE_ACTIVITY | 18 | -1.56 | 0.506 |
| 20 | NUCLEAR_DNA_DIRECTED_RNA_POLYMERASE_COMPLEX | 14 | -1.56 | 0.482 |

**Supplemental Table 3. Pathways upregulated based on RNA-seq analysis of the mock cell line after treatment with doxycycline**

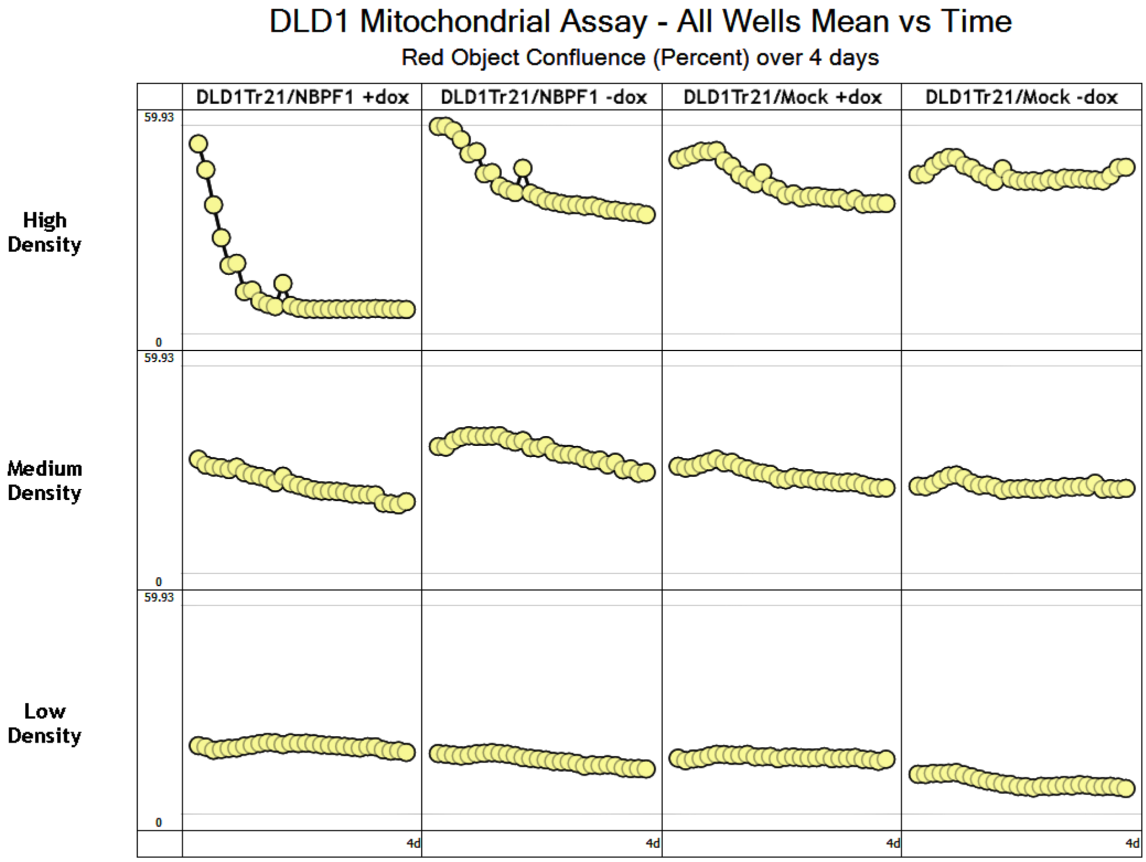


**Supplemental Figure 3. Red object confluence in different cell densities**

The immediate drop in red object confluence observed in *NBPF1*-induced cells appears to be dependent on cell density, as this is not observed at lower cell densities.

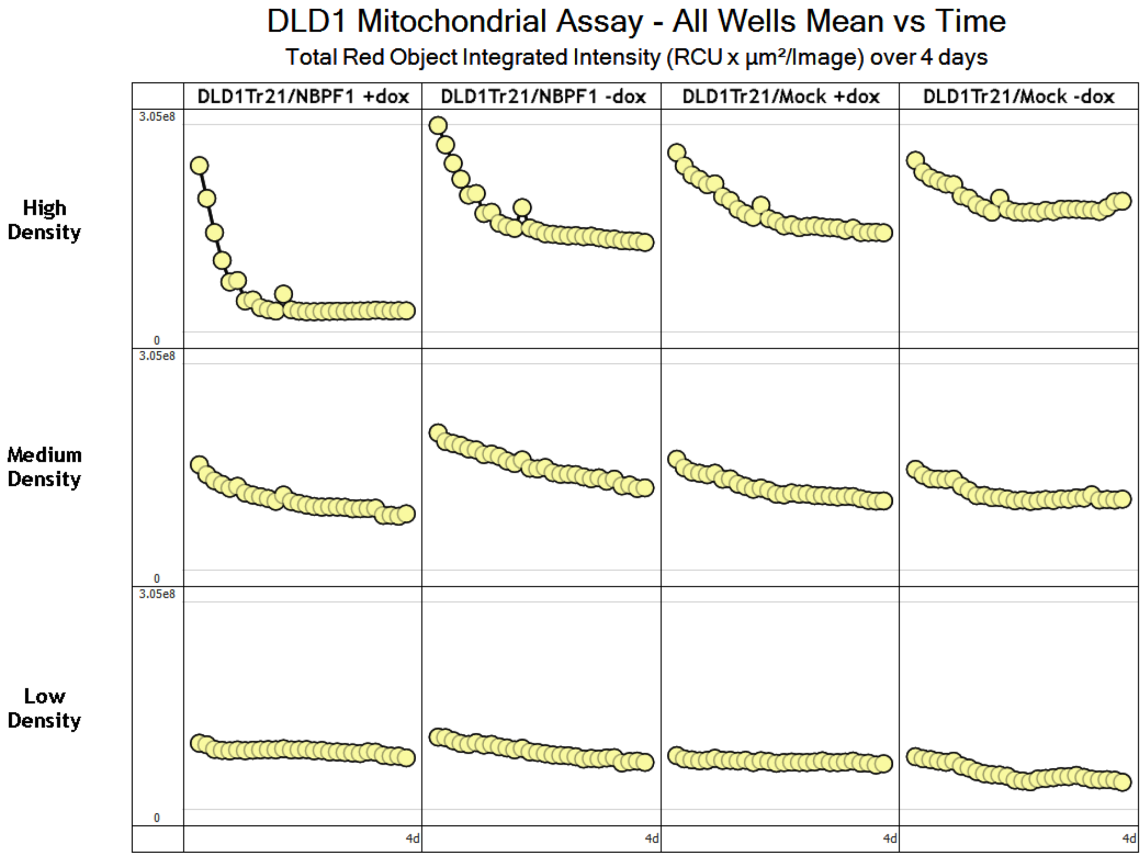


**Supplemental Figure 4. Red object intensity in different cell densities**

The suppression of red object intensity observed in *NBPF1*-induced cells is not present at lower cell densities.

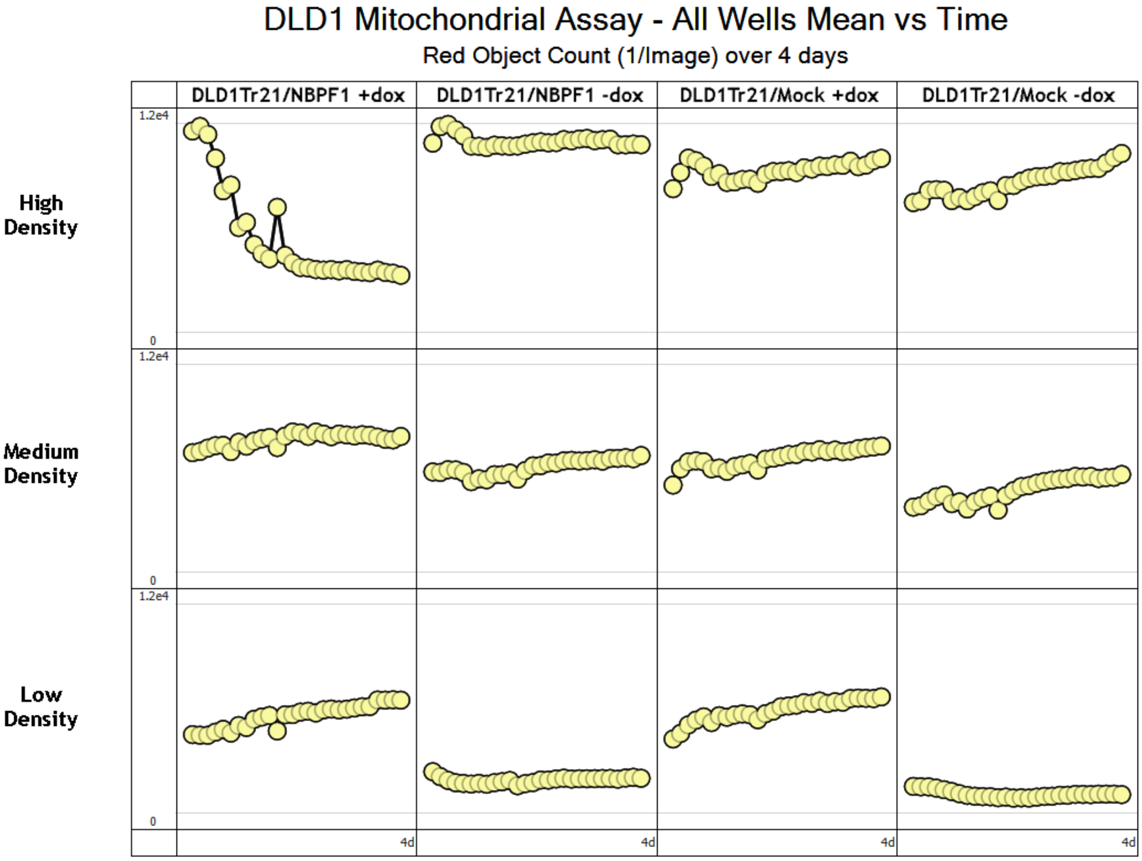


**Supplemental Figure 5. Red object count at different cell densities**

At high density, a reduction in the number of red objects is observed. At lower cell densities, the number of red objects measured is similar in *NBPF1*-induced cells to all control conditions.

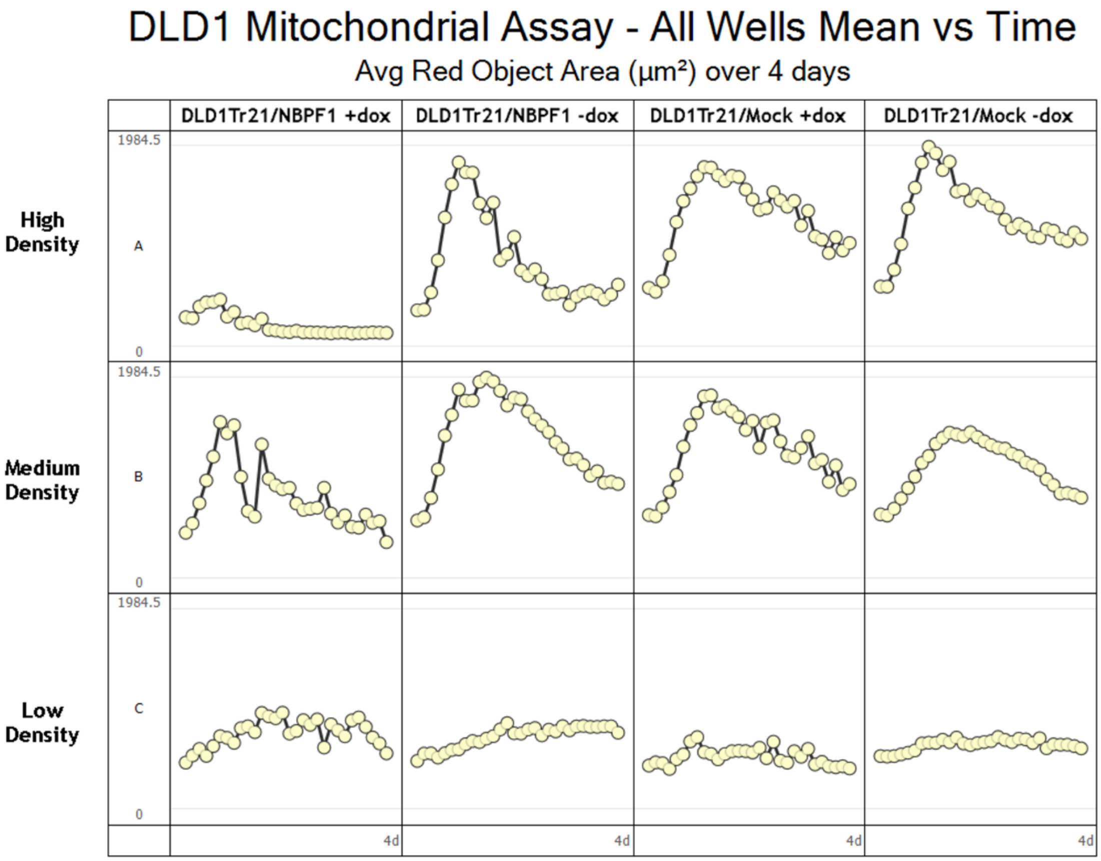


**Supplemental Figure 6. Average red object area by cell density**

The average size of red objects (in $\mu m^2$) when cell density is varied is shown. A substantial reduction in red signal is observed at high cell density and is not observed in doxycycline or *NBPF1* control conditions at any cell density.

| Sample | Expression Level | Doxycycline |
|---|---|---|
| sample_1 NBPF1 | 309.63 | + |
| sample_3 NBPF1 | 524.52 | + |
| sample_5 NBPF1 | 373.35 | + |
| sample_7 NBPF1 | 324.77 | + |
| sample_2 NBPF1 | 15.81 | - |
| sample_4 NBPF1 | 10.95 | - |
| sample_6 NBPF1 | 12.34 | - |
| sample_8 NBPF1 | 16.14 | - |
| sample_9 MOCK | 565.99 | + |
| sample_11 MOCK | 473.67 | + |
| sample_13 MOCK | 550.40 | + |
| sample_15 MOCK | 732.73 | + |
| sample_10 MOCK | 4.88 | - |
| sample_12 MOCK | 8.19 | - |
| sample_14 MOCK | 8.67 | - |
| sample_16 MOCK | 5.29 | - |

**Supplemental Table 4. EGFP FPKM expression values determined by including the exogenous EGFP construct sequence as part of the reference genome in the transcript mapping step**

Odd-numbered samples were induced with doxycycline and even-numbered samples were uninduced. Samples 1–8 represent the DLD1Tr21/NBPF1 cell line and samples 9–16 represent the DLD1Tr21/Mock cell line.

| GENE NAME | ACCESSION NUMBER | PROTEIN DESCRIPTION | PEPTIDES | FOLD CHANGE |
|---|---|---|---|---|
| ***EXOSC6*** | Q5RKV6 | EXOS6_HUMAN Exosome complex component MTR3 | 3 | 1.07 |
| ***NOP2*** | P46087 | NOP2_HUMAN Putative ribosomal RNA methyltransferase NOP2 | 9 | 1.14 |
| ***SLC25A6*** | P12236 | ADT3_HUMAN ADP/ATP translocase 3 | 13 | 1.37 |
| ***POLR1B*** | Q9H9Y6 | RPA2_HUMAN DNA-directed RNA polymerase I subunit RPA2 | 2 | 1.37 |
| ***ABCF2*** | Q9UG63 | ABCF2_HUMAN ATP-binding cassette sub-family F member 2 | 2 | 1.37 |
| ***DDX54*** | Q8TDD1 | DDX54_HUMAN ATP-dependent RNA helicase DDX54 | 3 | 1.39 |
| ***PQBP1*** | O60828 | PQBP1_HUMAN Polyglutamine-binding protein 1 | 4 | 1.39 |
| ***ATAD3A*** | Q9NVI7 | ATD3A_HUMAN ATPase family AAA domain-containing protein 3A | 9 | 1.39 |
| ***AFG3L2*** | Q9Y4W6 | AFG32_HUMAN AFG3-like protein 2 | 5 | 1.40 |
| ***RINT1*** | Q6NUQ1 | RINT1_HUMAN RAD50-interacting protein 1 | 1 | 1.40 |
| ***SLC25A10*** | Q9UBX3 | DIC_HUMAN Mitochondrial dicarboxylate carrier | 4 | 1.41 |
| ***KIAA0020*** | Q15397 | K0020_HUMAN Pumilio domain-containing protein KIAA0020 | 3 | 1.42 |
| ***MKI67*** | P46013 | KI67_HUMAN Antigen KI-67 | 6 | 1.43 |
| ***SMC2*** | O95347 | SMC2_HUMAN Structural maintenance of chromosomes protein 2 | 14 | 1.46 |
| ***SLC25A24*** | Q6NUK1 | SCMC1_HUMAN Calcium-binding mitochondrial carrier protein SCaMC-1 | 11 | 1.46 |
| ***XPO5*** | Q9HAV4 | XPO5_HUMAN Exportin-5 | 11 | 1.47 |
| ***COX17*** | Q14061 | COX17_HUMAN Cytochrome c oxidase copper chaperone | 1 | 1.48 |
| ***TOMM20*** | Q15388 | TOM20_HUMAN Mitochondrial import receptor subunit TOM20 homolog | 1 | 1.49 |

| | | | | |
|---|---|---|---|---|
| ***GLG1*** | Q92896 | GSLG1_HUMAN Golgi apparatus protein 1 | 2 | 1.49 |
| ***BAG3*** | O95817 | BAG3_HUMAN BAG family molecular chaperone regulator 3 | 3 | 1.51 |
| ***KPNA2*** | P52292 | IMA2_HUMAN Importin subunit alpha-2 | 8 | 1.55 |
| ***BCL2L13*** | Q9BXK5 | B2L13_HUMAN Bcl-2-like protein 13 | 5 | 1.55 |
| ***UQCRFS1*** | P47985 | UCRI_HUMAN Cytochrome b-c1 complex subunit Rieske, mitochondrial | 6 | 1.56 |
| ***C19ORF25*** | Q9UFG5 | CS025_HUMAN UPF0449 protein C19orf25 | 2 | 1.57 |
| ***SF3A2*** | Q15428 | SF3A2_HUMAN Splicing factor 3A subunit 2 | 3 | 1.58 |
| ***NDUFV2*** | P19404 | NDUV2_HUMAN NADH dehydrogenase [ubiquinone] flavoprotein 2, mitochondrial | 4 | 1.59 |
| ***SRSF11*** | Q05519 | SRS11_HUMAN Serine/arginine-rich splicing factor 11 | 3 | 1.60 |
| ***UQCRC1*** | P31930 | QCR1_HUMAN Cytochrome b-c1 complex subunit 1, mitochondrial | 10 | 1.60 |
| ***NDUFA5*** | Q16718 | NDUA5_HUMAN NADH dehydrogenase [ubiquinone] 1 alpha subcomplex subunit 5 | 4 | 1.62 |
| ***PHB*** | P35232 | PHB_HUMAN Prohibitin | 12 | 1.63 |
| ***MTCH2*** | Q9Y6C9 | MTCH2_HUMAN Mitochondrial carrier homolog 2 | 3 | 1.65 |
| ***IMMT*** | Q16891 | IMMT_HUMAN Mitochondrial inner membrane protein | 2 | 1.67 |
| ***VDAC1*** | P21796 | VDAC1_HUMAN Voltage-dependent anion-selective channel protein 1 | 5 | 1.69 |
| ***COX5B*** | P10606 | COX5B_HUMAN Cytochrome c oxidase subunit 5B, mitochondrial | 2 | 1.75 |
| ***YLPM1*** | P49750 | YLPM1_HUMAN YLP motif-containing protein 1 | 2 | 1.76 |
| ***PHB2*** | Q99623 | PHB2_HUMAN Prohibitin-2 | 11 | 1.76 |
| ***NDUFS2*** | O75306 | NDUS2_HUMAN NADH dehydrogenase | 5 | 1.79 |

| | | | | |
|---|---|---|---|---|
| | | [ubiquinone] iron-sulfur protein 2, mitochondrial | | |
| ***NDUFS8*** | O00217 | NDUS8_HUMAN NADH dehydrogenase [ubiquinone] iron-sulfur protein 8, mitochondrial | 3 | 1.80 |
| ***NDUFS1*** | P28331 | NDUS1_HUMAN NADH-ubiquinone oxidoreductase 75 kDa subunit, mitochondrial | 12 | 1.84 |
| ***NDUFS3*** | O75489 | NDUS3_HUMAN NADH dehydrogenase [ubiquinone] iron-sulfur protein 3, mitochondrial | 7 | 1.87 |
| ***STAU1*** | O95793 | STAU1_HUMAN Double-stranded RNA-binding protein Staufen homolog 1 | 10 | 1.96 |
| ***SLC25A13*** | Q9UJS0 | CMC2_HUMAN Calcium-binding mitochondrial carrier protein Aralar2 | 6 | 1.97 |
| ***VDAC2*** | P45880 | VDAC2_HUMAN Voltage-dependent anion-selective channel protein 2 | 4 | 2.00 |
| ***NDUFA10*** | O95299 | NDUAA_HUMAN NADH dehydrogenase [ubiquinone] 1 alpha subcomplex subunit 10, mitochondrial | 2 | 2.02 |
| ***UQCR C2*** | P22695 | QCR2_HUMAN Cytochrome b-c1 complex subunit 2, mitochondrial | 13 | 2.17 |
| ***TOMM22*** | Q9NS69 | TOM22_HUMAN Mitochondrial import receptor subunit TOM22 homolog | 3 | 2.34 |
| ***COX7A2*** | P14406 | CX7A2_HUMAN Cytochrome c oxidase subunit 7A2, mitochondrial | 2 | 2.42 |
| ***UQCRB*** | P14927 | QCR7_HUMAN Cytochrome b-c1 complex subunit 7 | 3 | 2.42 |
| ***S100P*** | P25815 | S100P_HUMAN Protein S100-P | 3 | 2.55 |
| ***MT-CO2*** | P00403 | COX2_HUMAN Cytochrome c oxidase subunit 2 | 2 | 2.56 |
| ***NDUFB4*** | O95168 | NDUB4_HUMAN NADH dehydrogenase [ubiquinone] 1 beta subcomplex subunit 4 | 2 | 3.72 |

| | | | | |
|---|---|---|---|---|
| ***KRT9*** | P35527 | K1C9_HUMAN Keratin, type I cytoskeletal 9 | 13 | 331.09 |

**Supplemental Table 5. List of downregulated protein candidates upon *NBPF1* expression**

At the 99% confidence interval, 79 differentially expressed proteins were identified. Gene name, accession number, protein description, peptide number, and fold change are shown here for 52 downregulated proteins.

| GENE NAME | ACCESSION NUMBER | PROTEIN DESCRIPTION | PEPTIDES | FOLD CHANGE |
|---|---|---|---|---|
| ***ARID1A*** | O14497 | ARI1A_HUMAN AT-rich interactive domain-containing protein 1A | 4 | 0.40 |
| ***PSMB9*** | P28065 | PSB9_HUMAN Proteasome subunit beta type-9 | 2 | 0.43 |
| ***GYG1*** | P46976 | GLYG_HUMAN Glycogenin-1 | 2 | 0.52 |
| ***PSMB10*** | P40306 | PSB10_HUMAN Proteasome subunit beta type-10 | 2 | 0.53 |
| ***SHROOM3*** | Q8TF72 | SHRM3_HUMAN Protein Shroom3 | 3 | 0.53 |
| ***TPM1*** | P09493 | TPM1_HUMAN Tropomyosin alpha-1 chain | 8 | 0.56 |
| ***PHGDH*** | O43175 | SERA_HUMAN D-3-phosphoglycerate dehydrogenase | 20 | 0.58 |
| ***ABHD14B*** | Q96IU4 | ABHEB_HUMAN Abhydrolase domain-containing protein 14B | 4 | 0.58 |
| ***CLIC4*** | Q9Y696 | CLIC4_HUMAN Chloride intracellular channel protein 4 | 12 | 0.59 |
| ***CSRP1*** | P21291 | CSRP1_HUMAN Cysteine and glycine-rich protein 1 | 4 | 0.59 |
| ***ASNS*** | P08243 | ASNS_HUMAN Asparagine synthetase [glutamine-hydrolyzing] | 16 | 0.60 |
| ***PSAT1*** | Q9Y617 | SERC_HUMAN Phosphoserine aminotransferase | 16 | 0.60 |
| ***ASS1*** | P00966 | ASSY_HUMAN Argininosuccinate synthase | 13 | 0.60 |
| ***UGT1A1*** | P22309 | UD11_HUMAN UDP-glucuronosyltransferase 1-1 | 9 | 0.61 |
| ***SERPINB6*** | P35237 | SPB6_HUMAN Serpin B6 | 5 | 0.61 |
| ***PPL*** | O60437 | PEPL_HUMAN Periplakin | 16 | 0.62 |
| ***CARS*** | P49589 | SYCC_HUMAN Cysteinyl-tRNA synthetase, cytoplasmic | 11 | 0.62 |
| ***FYCO1*** | Q9BQS8 | FYCO1_HUMAN FYVE and coiled-coil domain-containing protein 1 | 4 | 0.62 |

| | | | | |
|---|---|---|---|---|
| ***CHMP2A*** | O43633 | CHM2A_HUMAN Charged multivesicular body protein 2a | 2 | 0.63 |
| ***ALDH6A1*** | Q02252 | MMSA_HUMAN Methylmalonate-semialdehyde dehydrogenase [acylating], mitochondrial | 2 | 0.63 |
| ***SSSCA1*** | O60232 | SSA27_HUMAN Sjoegren syndrome/scleroderma autoantigen 1 | 3 | 0.63 |
| ***GSTM3*** | P21266 | GSTM3_HUMAN Glutathione S-transferase Mu 3 | 12 | 0.63 |
| ***MVP*** | Q14764 | MVP_HUMAN Major vault protein | 21 | 0.64 |
| ***ARHGAP18PE*** | Q8N392 | RHG18_HUMAN Rho GTPase-activating protein 18 | 5 | 0.64 |
| ***CHD1L*** | Q86WJ1 | CHD1L_HUMAN Chromodomain-helicase-DNA-binding protein 1-like | 3 | 0.64 |
| ***SCRIB*** | Q14160 | SCRIB_HUMAN Protein scribble homolog | 6 | 0.80 |
| ***HTATIP2*** | Q9BUP3 | HTAI2_HUMAN Oxidoreductase HTATIP2 | 7 | 0.94 |

**Supplemental Table 6. List of upregulated proteins after *NBPF* expression**

Little congruency is noted among listed proteins.

**Supplemental Table 7.** Table describing several physical and genetic characteristics of primates. Several characteristics of neurons are presented, including cortical neuron number, and genomic copy number of Olduvai domains. Table after Gould, 1977. Cortical neuron number from Roth and Dicke, 2005.

* 40 weeks is not a true reflection of full development, since postnatal growth continues for years.

| Primate | Gestation (weeks) | Complete hair covering | Carpal ossification centers at birth | 1st dentition (months) | 2nd dentition (years) | Growing period (years) | Life span (years) | Sexual maturity, female (years) | Number of cortical neurons (millions) | Olduvai Copy Number |
|---|---|---|---|---|---|---|---|---|---|---|
| Macaque | 24 | During gestation | All centers | 0.6-5.9 | 1.6-6.8 | 7 | 25 | — | 480 | ~30-35 |
| Gibbon | 30 | | 2-3 | 1.2-? | ?-8.5 | 9 | 33 | — | — | ~65-75 |
| Orangutan | 39 | Onset during | 2-3 | 4.0-13.0 | 3.5-9.8 | 11 | 30 | — | — | ~92 |
| Gorilla | 37 | gestation completed | — | 3.0-13.0 | 3.0-10.5 | 11 | 35 | 6-7 | 4,300 | ~99 |
| Chimpanzee | 34 | after birth | 2 | 2.7-12.3 | 2.9-10.2 | 11 | 35 | 9 | 6,200 | ~125 |
| Homo sapiens | 40* | Never completed | 0 | 6.0-24.0 | 6.0-20.0 | 20 | 70 | 13 | 11,500 | ~270-300 |
